# From methylation to myelination: epigenomic and transcriptomic profiling of chronic inactive demyelinated multiple sclerosis lesions

**DOI:** 10.1101/2023.01.12.523740

**Authors:** Assia Tiane, Melissa Schepers, Rick A. Reijnders, Lieve van Veggel, Sarah Chenine, Ben Rombaut, Emma Dempster, Catherine Verfaillie, Kobi Wasner, Anne Grünewald, Jos Prickaerts, Ehsan Pishva, Niels Hellings, Daniel van den Hove, Tim Vanmierlo

## Abstract

**Introduction:** In the progressive phase of multiple sclerosis (MS), the hampered differentiation capacity of oligodendrocyte precursor cells (OPCs) eventually results in remyelination failure. We have previously shown that DNA methylation of *Id2/Id4* is highly involved in OPC differentiation and remyelination. In this study, we took an unbiased approach by determining genome-wide DNA methylation patterns within chronically demyelinated MS lesions and investigated how certain epigenetic signatures relate to OPC differentiation capacity.

**Methods:** We compared genome-wide DNA methylation and transcriptional profiles between chronically demyelinated MS lesions and matched normal-appearing white matter (NAWM), making use of post-mortem brain tissue (n=9/group). DNA methylation differences that inversely correlated with mRNA expression of their corresponding genes were validated for their cell-type specificity in laser-captured OPCs using pyrosequencing. The CRISPR-dCas9-DNMT3a/TET1 system was used to epigenetically edit human-iPSC-derived oligodendrocytes to assess the effect on cellular differentiation.

**Results:** Our data show hypermethylation of CpGs within genes that cluster in gene ontologies related to myelination and axon ensheathment. Cell type-specific validation indicates a region-dependent hypermethylation of *MBP*, encoding for myelin basic protein, in OPCs obtained from white matter lesions compared to NAWM-derived OPCs. By altering the DNA methylation state of specific CpGs within the promotor region of *MBP*, using epigenetic editing, we show that cellular differentiation can be bidirectionally manipulated using the CRISPR-dCas9-DNMT3a/TET1 system *in vitro*.

**Conclusion:** Our data indicate that OPCs within chronically demyelinated MS lesions acquire an inhibitory phenotype, which translates into hypermethylation of crucial myelination related genes. Altering the epigenetic status of *MBP* can restore the differentiation capacity of OPCs and possibly boost (re)myelination.

## Introduction

Multiple sclerosis (MS) is a demyelinating disease of the central nervous system (CNS), characterised by a variety of clinical symptoms, such as visual problems, fatigue, muscle stiffness, and cognitive impairment (1). MS is defined by inflammation-induced demyelination during the early stages, which eventually results in gradual neurological disability as the disease progresses (1, 2).

During the progressive stages of MS, endogenous repair mechanisms (remyelination) become exhausted, resulting in the accumulation of chronically demyelinated lesions. Sustained demyelination within such lesions eventually causes loss of axonal density and neurodegeneration, which are two major contributors to the progressive nature of MS (3). Even though the exact aetiology of progressive MS remains unclear, it is suggested that remyelination is hampered in these stages due to the inability of oligodendrocyte precursor cells (OPCs) to differentiate into mature myelinating oligodendrocytes (4). Indeed, despite the abundant presence of OPCs within chronically demyelinated inactive MS lesions, their differentiation towards myelinating oligodendrocytes is attenuated in these demyelinated areas (5). This suggests that OPCs within chronically demyelinated MS lesions acquire a quiescent phenotype, leading to a differentiation block and, thus, ineffective remyelination.

In support of this idea, it has been shown that OPC differentiation is highly dependent on epigenetic regulation, which can be easily influenced by external stimuli from the surrounding microenvironment, such as sustained inflammation and inhibitory factors of the extracellular matrix (6–10). Epigenetic modifications are highly implicated in oligodendroglial biology (6, 11, 12). DNA methylation, for instance, is a stable yet at the same time dynamic epigenetic mark that translates environmental stimuli to alterations in gene expression and subsequent cellular behaviour. We and others have previously shown that DNA methylation contributes to physiological OPC differentiation (13, 14). In addition, DNA methylation is also required for remyelination, as shown in a mouse model for focal demyelination (15). This suggests that in the context of progressive MS, disturbed DNA methylation patterns in the oligodendrocyte lineage might be an acquired underlying feature of remyelination failure. Despite many advances in the field of neuroepigenetics, the number of epigenome-wide association studies (EWAS) conducted on MS brain tissue is very limited. The majority of EWAS studies in MS have been performed on normal-appearing white matter (NAWM) samples, which revealed important changes in DNA methylation prior to myelin damage, but do not show the epigenetic state in actual demyelinated MS lesions, where OPCs acquire a quiescent phenotype resulting in impaired remyelination (16–18).

In the present study, we quantified epigenomic and transcriptomic profiles of chronically demyelinated inactive MS lesions and their surrounding NAWM in order to investigate which genes could underlie the differentiation block of OPCs within the lesion environment. Cell-specific validation in laser-captured OPCs showed that OPCs within the lesion exhibit a hypermethylated profile of essential myelin genes, such as *MBP*. By applying the CRISPR/dCas9-mediated epigenetic editing toolbox, we examined the causal relationship between the methylation of these myelin genes and the differentiation capacity of human induced pluripotent stem cell (iPSC)-derived oligodendrocytes.

## Materials and methods

### Sample collection

Human post-mortem brain tissue was obtained from the Netherlands Brain Bank (www.brainbank.nl). Chronic, inactive demyelinated white matter lesions were selected from progressive MS patients (n=10, one lesion per patient) and characterised for demyelination (Proteolipid protein [PLP^-^]), inflammation (Human leukocyte antigen [HLA-DR^-^], Oil Red O [ORO^-^]), and presence of OPCs (Neural/glial antigen 2 [NG2^+^]) by immunohistochemistry (13). Demographic details were described before (13). Lesions were manually dissected from the surrounding NAWM, using the proteolipid protein (PLP) staining as a reference. Slices of 30 μM were made using a CM3050 S cryostat (Leica) and were alternately collected for either RNA or DNA isolation (Figure 1a). For laser-capture microdissection and immunohistochemistry, slices of 10 μM were cut, and collected on glass microscopy slides.

**Fig. 1.**
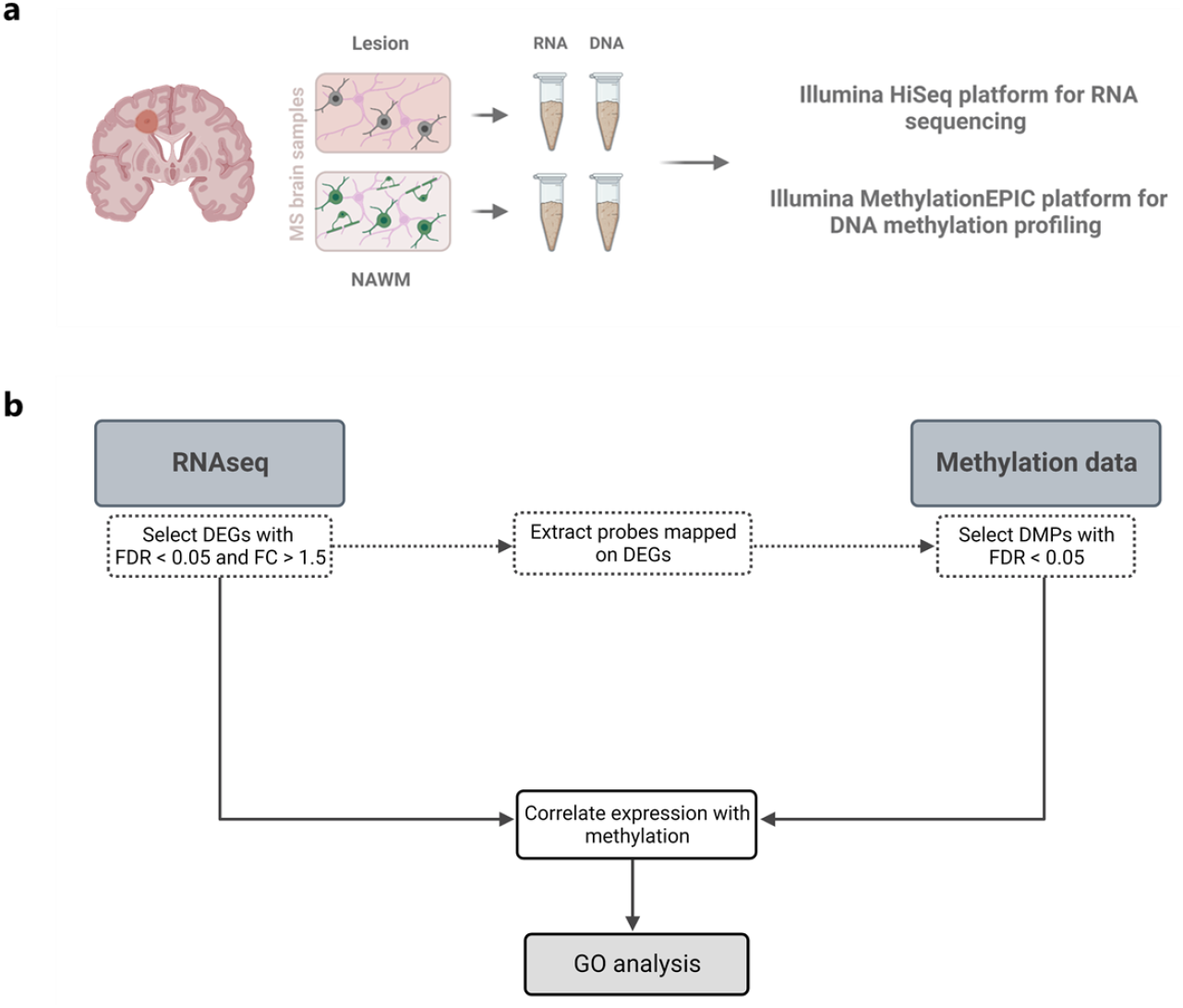
Overview of the sample preparation and data analysis workflow. **a** Multiple sclerosis (MS) lesions and surrounding normal appearing white matter (NAWM) were dissected and both were collected for RNA and DNA isolation. Transcriptomic and methylomic profiling was carried out using the HISeq sequencing and Illumina MethylationEPIC array platform, respectively. **b** Illustration of the data analysis workflow integrating the transcriptomic and methylomic datasets. NAWM: normal-appearing white matter, DEGs: differential expressed genes, FDR: false discovery rate adjusted p-value, FC: fold change, DMPs: differential methylated probes, GO: gene ontology.

### Transcriptomic profiling

Total RNA was extracted from lesions and their surrounding NAWM, using the RNeasy mini kit (Qiagen), according to the manufacturer’s instructions. RNA concentrations were analysed with a Nanodrop spectrophotometer (Isogen Life Science). RNA integrity was checked using the Agilent RNA 6000 Pico Bioanalyzer (Agilent Technologies). RNA integrity number (RIN) values ranged between 2.40 and 6.70. Samples were processed and sequenced by the Genomics Core Leuven (Leuven, Belgium). Library preparation was performed using the Lexogen 3’mRNA-Seq Library Prep Kit (Isogen Life Science). Libraries were sequenced on the Illumina HiSeq4000 sequencing system. Quality control (QC) of raw reads was performed with FastQC v0.11.7 (19). Adapters were filtered with ea-utils fastq-mcf v1.05 (20). Using the default parameters, splice-aware alignment was performed with HISat2 against the human reference genome hg38 (21). Reads mapping to multiple loci in the reference genome were discarded. The resulting BAM alignment files were handled with Samtools v1.5 (22). Read counts for each gene were compiled using Rsubread (version 2.8.2) by reading in and processing each bam file. A minimum threshold of 15 counts per million reads for at least 40% of all samples was used to determine whether a gene was expressed, leaving 8399 genes for analysis. The package EdgeR (version 3.36.0) was used to normalise and transform counts to log counts-per-million, using the Trimmed Mean of M-values (TMM) normalisation method.

### Methylomic profiling

Genomic DNA was extracted using a standard chloroform-phenol extraction and ethanol precipitation method. DNA concentration was assessed with the Qubit dsDNA HS Assay Kit (Invitrogen). A minimum of 500 ng per sample was used for the Illumina Infinium MethylationEPIC array BeadChip (850K), which was carried out by the Epigenomic Services from Diagenode (Liège, Belgium; Cat nr. G02090000). The DNA was deaminated with the EZ-96 DNA Methylation Kit (Zymo Research) according to Illumina’s recommended deamination protocol. Methylation data processing and statistical analyses were performed using the programming language R (version 4.1.2.) and RStudio (version 2021.09.1). Raw IDAT files were loaded into R using the minfi package (23). To confirm that matched lesion and NAWM samples were from the same individual, we made use of 59 single nucleotide polymorphism (SNP) probes on the Illumina EPIC array to cluster genetically identical samples. Cell proportion estimates were generated using the Houseman method (24). Samples with a NeuN^+^ estimation of more than 5% were excluded from the analysis. Cross-hybridizing probes and probes containing SNPs were removed (25). Probe filtering was performed using the *pfilter* function from the wateRmelon package (version 2.0.0) to exclude probes with >1% of samples with a detection p-value >0.05 (26). The remaining data were normalised using the *dasen* function from the wateRmelon package, and probes on the X and Y chromosomes were excluded from the dataset. As principle component analysis (PCA) trait analysis showed a significant correlation with the EPIC chip IDs, we corrected for this batch effect using the *ComBat* function from the *sva* package (version 3.20.0), which applies a Bayesian method to adjust for known batch covariates (27). After data processing, eight lesion and nine NAWM samples remained, as well as 769,804 probes.

### Laser-capture microdissection

OPCs were stained using an accelerated protocol to maintain DNA integrity. Briefly, sections were fixed in ice-cold acetone for 10 minutes and dip-washed in TBS/TBS-T/TBS. Endogenous peroxidase activity was neutralised with 1.5% H_2_O_2_ in TBS for 10 seconds, followed by a rinse with TBS and a 30-minute blocking step with the Dako Protein Block (Dako) at room temperature. The slices were incubated with a primary antibody against NG2 (1:200, Abcam Ab101807) for 30 minutes, followed by a quick wash step in TBS. Sections were incubated with horseradish peroxidase (HRP)-conjugated EnVision + Dual Link System (Dako) for 15 minutes, washed with TBS and incubated with an avidin-biotinylated horseradish peroxidase complex for 10 minutes, after which visualisation of the staining was accomplished using 0.3% ammonium nickel sulphate and 0.025% diaminobenzidine (pH 7.8) in TBS. After sequential dehydration steps (30 seconds in 75%-95%-100% ethanol and five minutes in xylene), the samples were ready for immediate laser-capture microdissection using a PALM MicroBeam (Zeiss). 50 cells were isolated per region and collected into 0.1 ml tube caps containing 10 μl PBS.

### CRISPR-dCas9 plasmids

#### Guide design

A specific single guide RNA (sgRNA) was designed to induce (de)methylation within the promoter region of the *MBP* (chr18:74,690,791-74,691,721) gene using Benchling software (Supplementary Table 1). Guides were synthesised as oligos with overhangs to fit into the BbsI restriction gap and an additional guanine for increased transcriptional efficiency.

#### sgRNA cloning

Guides were cloned into the DNMT3a plasmids (Addgene #71667 and #71684) using a one-step digestion and ligation protocol. Briefly, 100 ng of plasmid was added to a mixture of 1 μM of the annealed guide oligos, 20 U BbsI restriction enzyme (Bioké), 1x cutsmart buffer (Bioké), 400 U T4 ligase (Bioké), 1x T4 ligase buffer (Bioké) and H_2_O to an end volume of 20 μl and incubated for 30 cycles of 5 minutes on 37°C and 5 minutes on 23°C. The product was then transformed into NEB 5-alpha Competent E. coli cells (Bioké) and plated on LB-agar plates, supplemented with ampicillin (Amp; 100 mg/ml). Suitable colonies were propagated overnight in LB-Amp medium. Plasmids were extracted using the NucleoBond Xtra Midi kit, according to the manufacturer’s protocol (Macherey-Nagel). Sanger sequencing was carried out on purified plasmid vector to validate the sgRNA incorporation. For the TET1 vectors (Addgene #129025 and #129026), we performed subcloning from the DNMT3a vectors using the PvuI and XbaI restriction enzymes (Thermofisher). One μg of each vector was incubated overnight at 37°C with 10 U of both restriction enzymes, 1x Tango buffer, and H2O up to a total volume of 50 μl. The samples were loaded on an agarose gel (1%) and both insert (from the DNMT3a vectors), as well as vectors (from the TET1 vectors) were extracted from the gel, using the PCR and gel clean-up kit (Macherey-Nagel) according to the manufacturer’s instructions. Inserts and vectors were ligated with the T4 DNA Ligase buffer and enzyme system (Bioké) into the linearized vector in a 2:1 insert to vector molar ratio. Plasmid transformation and purification was performed as described above.

### Cell culture and transfection

#### Human-derived iPSC-oligodendrocytes

Inducible SOX10-overexpressing iPSCs were used to generate O4+ and MBP+ oligodendrocyte cultures as described previously and kindly provided under a material transfer agreement (MTA) by Catherine Verfaillie (KuLeuven, Leuven, Belgium) (28, 29). Differentiated iPSC-oligos were frozen in liquid nitrogen and thawed for transfection experiments. Cells were seeded at a density of 250,000 cells/well in a PLO/laminin-coated 24-well plate and maintained in differentiation medium with doxycycline (4 μg/ml). The DNMT3a plasmids were a gift from Vlatka Zoldoš (Addgene #71667 and #71684), and the TET1 plasmids were a gift from Julia K. Polansky (Addgene #129025 and #129026). Plasmids were transfected into human iPSC-derived OPCs 48 hours after seeding, using the OZ Biosciences NeuroMag Transfection Reagent (Bio-connect), following the manufacturer’s instructions. In brief, 1 μg of plasmid DNA was diluted in 50 μl DMEM/F12 medium, added to 1.75 μl NeuroMag reagent and incubated for 20 minutes at room temperature. DNA/NeuroMag complexes were dropwise added to iPSC-oligo cultures (250,000 cells/well), maintained in differentiation medium, and placed on a magnetic plate for 4 hours in a 5% CO2 incubator. Medium change with fresh differentiation medium, containing doxycycline (4 μg/ml), was performed 72 hours after transfection. Cells were lysed or fixated on day five post-transfection for further experiments.

#### Human oligodendroglioma (HOG) cell line

The human oligodendroglioma cell line HOG was maintained in culture medium (DMEM, 10% FCS, 1% P/S) at 37°C and 5% CO2. For transfection experiments, cells were seeded in poly-L-lysine (PLL, Sigma-Aldrich)-coated 24-well plates at a density of 37,500 cells per well. After attaching to the plate, cells were transfected on the same day as the seeding, using the protocol described above, with a minor adjustment (3 μl NeuroMag reagent and 500 ng DNA per well for 30 minutes on the magnetic plate). Cells were maintained in differentiation medium (DMEM, 1% P/S, 0.05% FCS, 5 μg/ml bovine insulin, 5 μg/ml transferrin, 0.03 nM sodium selenite, 30 nM L-thyroxine; all from Sigma-Aldrich), with one medium change 48 hours after transfection. On day four post-transfection, cells were fixated for further experiments.

### Pyrosequencing

Genomic DNA was extracted from laser-captured OPCs, as well as transfected iPSC-OPCs, and bisulfite-converted, using the Zymo Research EZ DNA Methylation-Direct Kit (BaseClear Lab Products). PCR primers were designed using the PyroMark Assay Design 2.0 software (Qiagen, Supplementary Table 2). The assay for *MBP* was tested for its sensitivity using the EpiTect PCR Control DNA Set (Qiagen). Product amplification was performed using the following reaction mixture: 1X Buffer with 20 mM MgCl2 (Roche), 10 mM dNTP mix (Roche), 5 μM forward and reverse primers (Metabion AG), 1U FastStart Taq DNA Polymerase (Roche), bisulfite-converted DNA and nuclease-free water to a total volume of 25 μl. PCR cycling was performed as follows: initial denaturation for 5 minutes at 95 °C, 50 cycles of 30 seconds at 95 °C, 30 seconds at 60 °C and 1 minute at 72 °C; final extension for 7 minutes at 72 °C. PCR amplicons were sequenced using the Pyromark Q48 instrument (Qiagen) with the PyroMark Q48 Advanced CpG Reagents (Qiagen), according to the manufacturer’s protocol and quantified with the Pyromark Q48 Autoprep software.

### Quantitative PCR

Transfected iPSC-OPCs and post-mortem human MS samples were lysed in Qiazol (Qiagen), and RNA was isolated using a standard chloroform extraction and ethanol precipitation method. RNA concentration and quality were analysed with a Nanodrop spectrophotometer (Isogen Life Science). RNA was reverse-transcribed using the qScript cDNA Supermix kit (Quanta). qPCR was performed to analyse gene expression using the Applied Biosystems QuantStudio 3 Real-Time PCR System (Life Technologies). The reaction mixture consisted of SYBR Green master mix (Life Technologies), 10μM forward and reverse primers (Integrated DNA Technologies), nuclease-free water and cDNA template (12.5 ng), up to a total reaction volume of 10 μl. The primers used for amplification are listed in Supplementary Table 3. Start fluorescence values were calculated for the human MS sample validation of the RNAseq data. Transfection results were analysed by the comparative Ct method and were normalised to the most stable housekeeping genes (RPL13a and TBP).

### Immunocytochemistry

Transfected cells were fixed in 4% paraformaldehyde (PFA) for 30 minutes at room temperature. Aspecific binding was blocked for 30 minutes with 1% bovine serum albumin (BSA) in 0.1% PBS-T, followed by incubation with primary antibodies (Supplementary Table 4) for four hours at room temperature. After three washing steps with PBS, cells were incubated with Alexa 488- or Alexa 555-conjugated secondary antibody (Supplementary Table 4) for one hour. Nuclei were counterstained with 4’6-diamidino-2-phenylindole (DAPI; Sigma-Aldrich). Coverslips were mounted with Dako mounting medium (Dako) and analysed using a fluorescence microscope (Leica DM2000 LED). Per coverslip, three images were quantified using Fiji ImageJ software.

### Statistical analysis

Differential expression analysis was performed using the *limma* package (version 3.50.0) (30). Age, sex, and post-mortem interval (PMI) were included as covariates and individual was treated as a random intercept, using the *duplicateCorrelation* function from the *limma* package. P-values were FDR-corrected for multiple testing to determine differentially expressed genes (DEG, FDR p-value < 0.05 and absolute fold change >1.5) between lesion and NAWM samples.

We extracted all the CpG probes from the Illumina methylationEPIC array that were annotated (Illumina UCSC annotation) to DEGs from the RNAseq analysis. Out of the 769,804 probes, 29,446 probes were used as input for the differential methylation analysis using the same approach as for the DEG analysis. The *duplicateCorrelation* function from the *limma* package was applied to block individual as a random effect. Age, sex, and PMI were included as covariates in the regression model and FDR correction for multiple testing was applied to the nominal p-values to identify differentially methylated probes (DMPs, FDR p-value < 0.05).

All DE genes that contained a DMP were subjected to a Pearson’s correlation analysis between expression (LogCPM) and methylation (bèta values) levels. A final list of genes that displayed differential expression and methylation, and a significant correlation between both expression and methylation, was used as input for a gene ontology (GO) analysis using the *enrichGO* function from the *clusterProfiler* package (version 4.2.2), focusing on the ‘Biological Process’ ontologies. An overview of the data analysis workflow is provided in Figure 1b.

Statistical analysis of the transfection and pyrosequencing experiments was performed using GraphPad Prism 9.0.0 software (GraphPad software Inc., CA, USA). Differences between group means were determined using an unpaired t-test for normally distributed data and a Mann-Whitney test for not normally distributed data. Differences in methylation at different CpG sites were determined using a two-way repeated measures ANOVA with Šídák’s multiple comparisons test. All data are depicted as mean ±SEM, *=p≤0.05, **=p<0.01, ***=p<0.001, ****=p<0.0001.

## Results

### Transcriptomic profiling of chronic inactive demyelinated MS lesions and the surrounding NAWM

Bulk RNA sequencing was performed on chronically demyelinated MS lesions and the corresponding surrounding NAWM. After stringent data pre-processing and QC filtering, 17 samples (9 lesions and 8 NAWM) were included in the RNA data analysis. Gene clustering based on absolute expression levels indicated clustering of the lesions separately from the NAWM (Fig. 2a). Interestingly, lesions and NAWM that cluster close to each other were derived from the same individual. PCA based on the logCPM values showed that 63% of the variance could be explained by PC1, which highly correlated (p = 0.00059) with the sample group (Fig. 2b). Out of the total of 8,399 genes that were subjected to a differential gene expression analysis, 641 genes were found to be significantly differentially expressed between lesion and NAWM, with an absolute fold change above 1.5 (Fig. 2c, Supplementary Data 1). Interestingly, the distribution was roughly balanced between upregulated (242) and downregulated (399) genes.

**Fig. 2.**
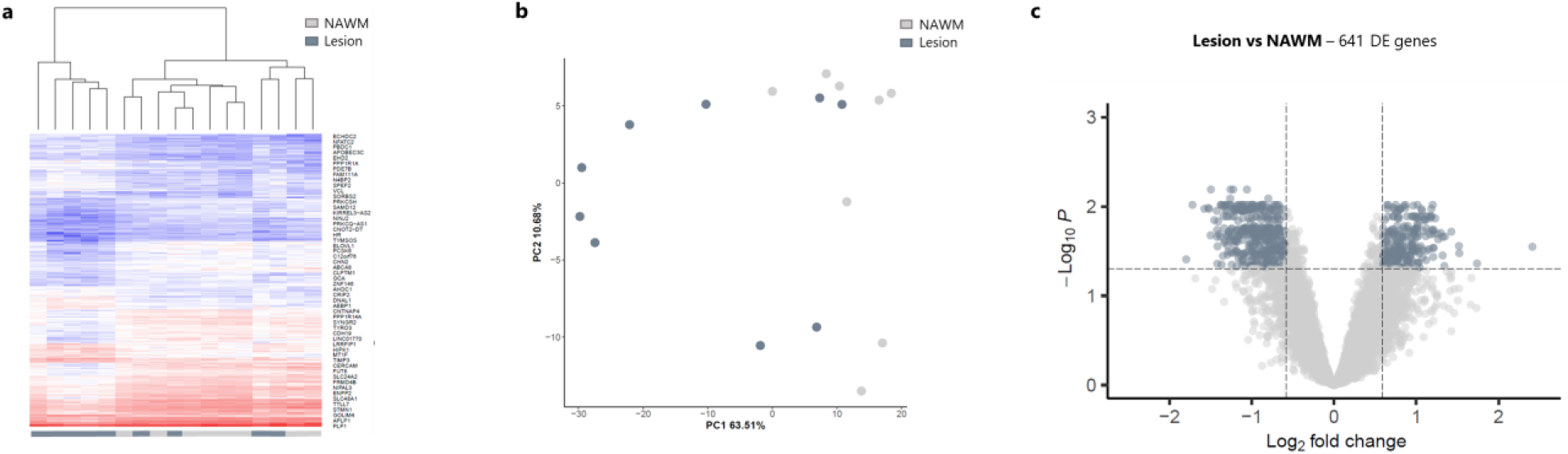
Chronically demyelinated lesions are transcriptionally distinct from the surrounding normal appearing white matter (NAWM). Based on the transcriptomic profile, chronic multiple sclerosis (MS) lesions can be distinguished from the surrounding NAWM, as determined by (**a**) gene clustering based on absolute expression levels and (**b**) a principal component analysis (PCA). **c** Differential expressed genes (DEGs) analysis revealed 641 genes that are significantly differentially expressed between lesion and NAWM (FDR p-value < 0.05), with an absolute fold change above 1.5.

### Genes involved in glial cell development and myelination are differentially methylated in chronic MS lesions

An EWAS was conducted using the Illumina methylationEPIC array to analyse the DNA methylation state of the chronically demyelinated lesions and NAWM samples. PCA revealed clustering of the samples based on the methylation bèta values, similar to those observed in the RNA sequencing data (Fig. 3a). Out of the 769,804 CpGs that passed the initial quality control, 29,446 CpG sites were annotated to the DEGs from the transcriptome analysis. Differential methylation analysis of these genes showed that 8,336 CpG positions were significantly (FDR p-value < 0.05) differentially methylated between lesions and NAWM (Fig. 3b, Supplementary Data 2). These differentially methylated positions (DMPs) were then subjected to a correlation analysis with the matching expression data. Interestingly, 508 genes showed a significant (FDR-adjusted p-value < 0.05) and strong correlation between their expression and methylation profile (Supplementary Data 3). Fig. 3c. shows the top ten correlating CpGs, nine of which showing a strong negative correlation between DNA methylation and RNA expression. The final set of 512 genes, which were differentially expressed, differentially methylated and correlated between both expression and methylation, was used for the GO analysis, with a focus on Biological Process (Fig. 3d). Clustering of the significantly enriched GO terms showed two main clusters, related to glial cell development/myelination and cytoskeleton organisation (Fig. 3d).

**Fig. 3.**
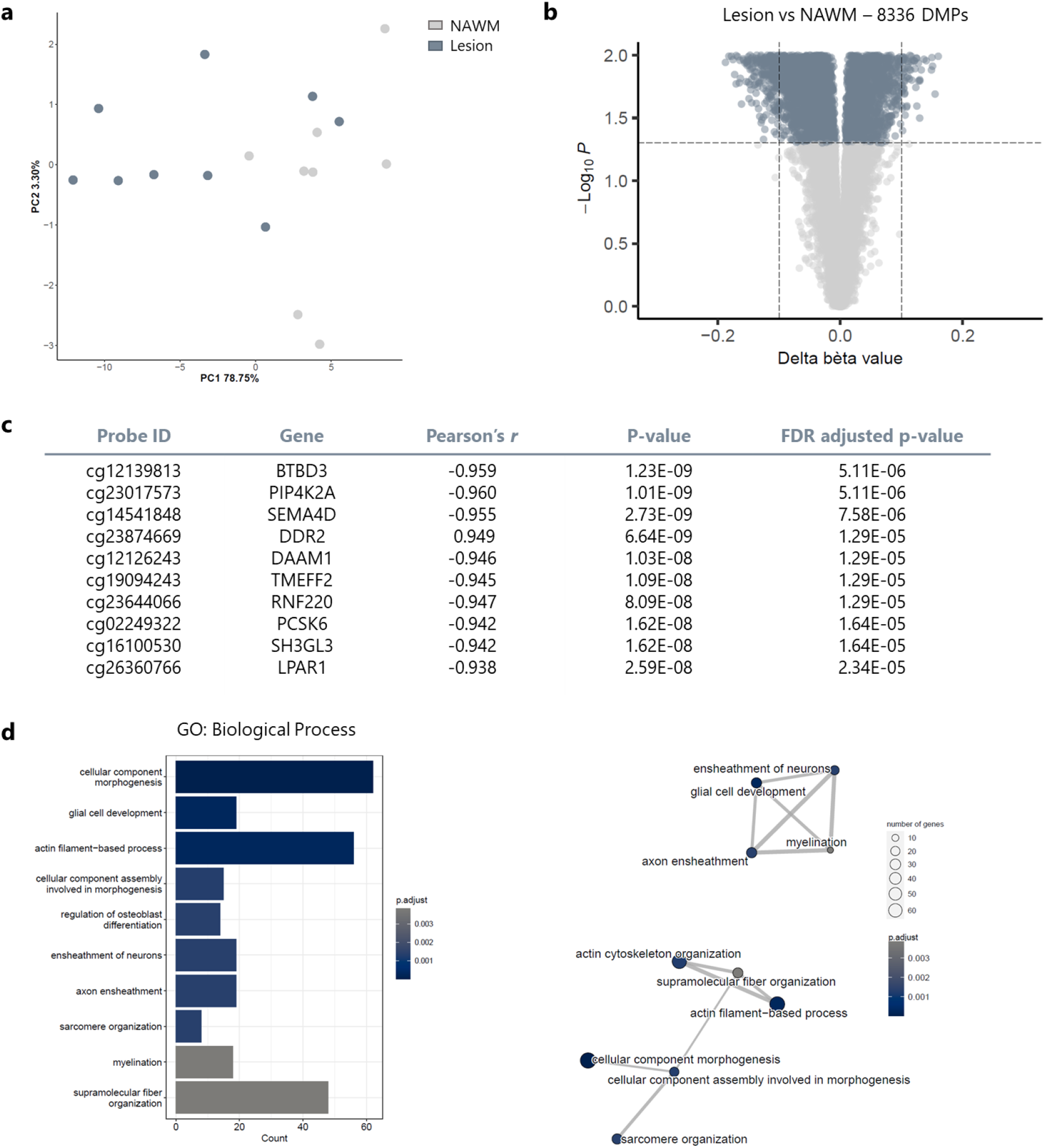
Differentially methylation analysis between lesions and NAWM reveals enriched gene ontologies (GOs) related to glial cell development and myelination. **a** Principal component analysis (PCA) shows the clustering of the samples based on the methylation bèta values. **b** Out of the 29,446 analysed CpG sites, 8,336 CpGs are differentially methylated between lesions and normal appearing white matter (NAWM; FDR<0.05). **c** Pearson’s correlation analysis between methylation and expression levels of the significantly differentially methylated CpGs. **d** Gene ontology analysis of the 512 genes that were differentially expressed and correlated significantly with their differential methylated probes (DMPs) revealed two main significantly enriched clusters related to the cytoskeleton and glial cell development/myelination.

As we are particularly interested in the contribution of DNA methylation to (re)myelination in the MS lesions, we focused on the genes that were part of the enriched GO clusters related to glial cell development/myelination (GO:0021782, GO:0007272, GO:0008366, GO:0042552). We explored the distribution of those DMPs across gene features (Fig. 4a) and CpG-related island features (Fig. 4b). Interestingly, the gene *MBP*, coding for myelin basic protein, the second most abundant protein in central nervous system myelin, did contain the highest number of DMPs in general as well as the highest number of DMPs that were located in the promotor region (TSS1500, TSS200) (Fig. 4a). An essential portion of these DMPs was furthermore situated in CpG islands or shores (Fig. 4b). Interestingly, all the CpGs within the promotor region of the gene were consistently hypermethylated in lesions compared to the surrounding NAWM (Fig. 4c). To technically validate our findings from the RNAseq and EWAS data, we performed targeted analysis of the expression and methylation profile of *MBP* using qPCR and pyrosequencing, respectively. The correlation analyses for both expression and methylation showed a strong and significant correlation between the two techniques, serving as a robust validation of the RNA sequencing and EWAS discovery data (Fig. 4d). Furthermore, we observed a significant negative correlation between *MBP* expression and methylation levels (Fig.4e). Altogether, these data suggest an important role of DNA methylation for the regulation of *MBP* expression.

**Fig. 4.**
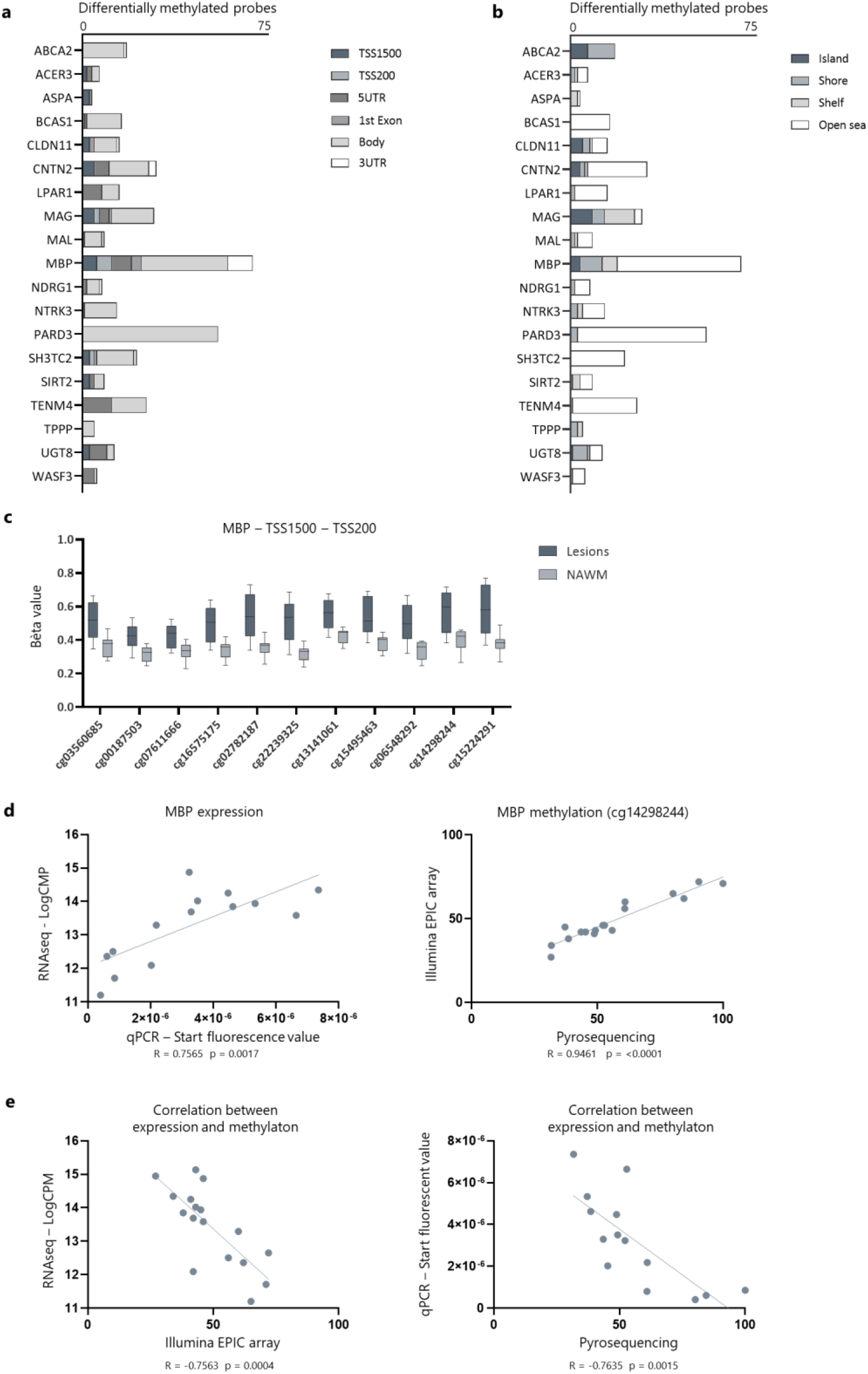
In-depth overview of the genes that are part of the enriched GO clusters related to glial cell development/myelination (GO:0021782, GO:0007272, GO:0008366, GO:0042552) Distribution of DMPs within the GO clusters related to myelination across gene features (**a**)and CpG-related island features (**b**). The height of the bars represents the number of DMPs annotated to the gene. **c** The beta values of the DMPs located in the promoter region (TSS1500, TSS200) of *MBP* indicate consistent hypermethylation within lesions compared to the surrounding NAWM. **d** Technical validation of the expression and methylation levels of MBP, as determined by qPCR and pyrosequencing, respectively. Pearson’s correlation analysis shows a significant correlation between both techniques on the expression level, as well as DNA methylation level (n=14). **e** Expression and methylation levels of *MBP* are significantly negatively correlated (Pearson’s correlation analysis), both for array-based techniques (RNAseq, Illumina EPIC array), as well as targeted techniques (qPCR, pyrosequencing) (n=14).

### Cell type-specific validation indicates hypermethylation of *MBP* in OPCs obtained from lesions, compared to NAWM-derived OPCs

The methylation signature within MS lesions suggests a potential differentiation and (re)myelination block, directly acting on essential myelin genes, such as *MBP*. However, as the Illumina EPIC array was performed on bulk tissue, the observed degree of methylation of *MBP* could also be explained by cellular heterogeneity of the samples. As we were particularly interested in whether there is a contribution of OPCs to the observed epigenetic signature of *MBP*, we stained OPCs within the samples, laser capture micro-dissected, and collected them for targeted methylation analysis of the *MBP* promotor region by means of pyrosequencing (Fig. 5a). In line with our bulk tissue findings, we again observed a hypermethylated profile in OPCs obtained from lesions compared to OPCs that were located in the NAWM (Fig. 5b, c).

**Fig. 5.**
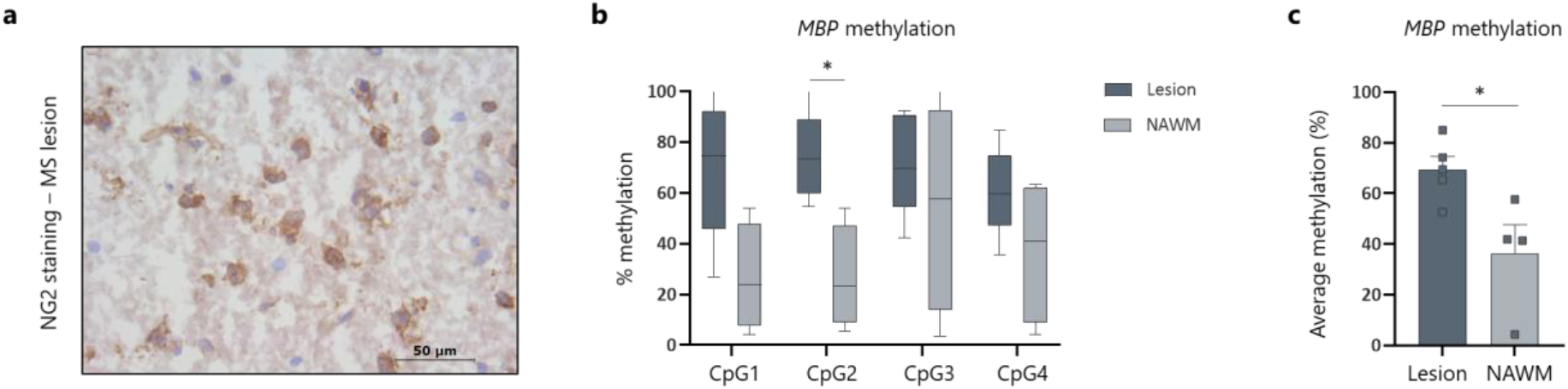
Cell-specific validation of the hypermethylated profile of *MBP* within OPCs derived from lesions, compared to OPCs isolated from the surrounding NAWM. **a** OPCs were stained for the NG2 marker and laser-capture microdissected from either lesions of NAWM. Batches of 50 cells per sample were subjected to bisulfite pyrosequencing to determine the methylation profile of the *MBP* promoter region. **b, c** OPCs within the promoter region of *MBP* show a hypermethylated profile compared to OPCs isolated from the NAWM (n=4-5, two-way ANOVA with Šídák’s multiple comparisons test for b and Wilcoxon test for c). Data are represented as mean ±SEM, *p<0.05.

### Targeted epigenetic editing of the *MBP* gene influences the differentiation capacity of human iPSC-derived oligodendrocytes and human oligodendroglioma cells

As we discussed elaborately in our recently published perspective (31), most EWAS observations remain correlational, making it difficult to infer a cause-effect relationship. In the last couple of years, epigenetic editing, i.e., altering the epigenome by directing e.g. DNA methylation at a specific site, has grown as a powerful tool to further study the role of epigenetics in health and disease, especially in view of addressing causality. Hence, we applied CRISPR-dCas9-based epigenetic editing to investigate potential cause-and-effect relationships for epigenetic alterations of *MBP* regarding oligodendrocyte differentiation. A sgRNA was designed to target the promotor region of *MBP* and cloned into CRISPR-dCas9-DNMT3a or CRISPR-dCas9-TET1 vectors, to respectively methylate or demethylate the CpG sites within the *MBP* promotor region. As the off-target effects, due to unspecific binding of the sgRNAs, still remain an ongoing challenge in the field of epigenetic editing, we used an online tool to predict possible off-target site of our sgRNA. All the predicted off-target sites were located in non-coding genomic regions of the DNA (Supplementary Table S5). We then explored the impact of epigenetic editing on human iPSC-derived oligodendrocyte differentiation. As MBP is a solid marker for mature oligodendrocytes, we performed immunostaining for MBP to observe the effects on the protein level, as well as to visualise and assess the cellular morphology (Fig. 6a, b). Cells transfected with an active TET1 construct targeting the *MBP* promotor showed increased MBP protein expression (as determined by the MBP-positive area per transfected cell). Human iPSC-derived oligodendrocytes that were transfected with the DNMT3a construct to methylate the *MBP* promotor showed a tendency towards decreased MBP expression. To evaluate the differentiation capacity of the transfected cells, we furthermore performed a Sholl analysis (Fig. 6c-f). Analysis of the ending radius (Fig. 6d), the sum of intersections (Fig. 6e) and the average number of intersections per Sholl ring (Fig. 6f) all showed that modulation of the *MBP* promotor methylation status influences cellular differentiation. Interestingly, we observe an overall more pronounced effect in the TET1-mediated demethylation experiments compared to DNMT3a-driven targeted methylation. In line, we observed a trend towards lower methylation levels and higher expression levels of *MBP* after targeted demethylation (Fig. 6g-h). Next to the low statistical power, our heterogeneous bulk cultures consisted of both transfected as well as untransfected cells, leaving our expression and methylation results confounded by the background noise of unmodified cells. Off note, the functional readouts, which were based on transfected cells only, showed strong and significant results after epigenetic editing of *MBP* (Fig. 6a-f). To validate our findings, we also transfected a human oligodendroglioma cell line with the epigenetic editing vectors (Fig. 6i). Similar effects were observed as to iPSC-derived oligodendrocytes, both in terms of cellular complexity (Fig. 6j) and MBP fluorescence area (Fig. 6k) of the transfected cells. Altogether, these results show that by altering the methylation profile of the *MBP* gene, it is possible to influence the differentiation capacity of human oligodendrocytes.

**Fig. 6.**
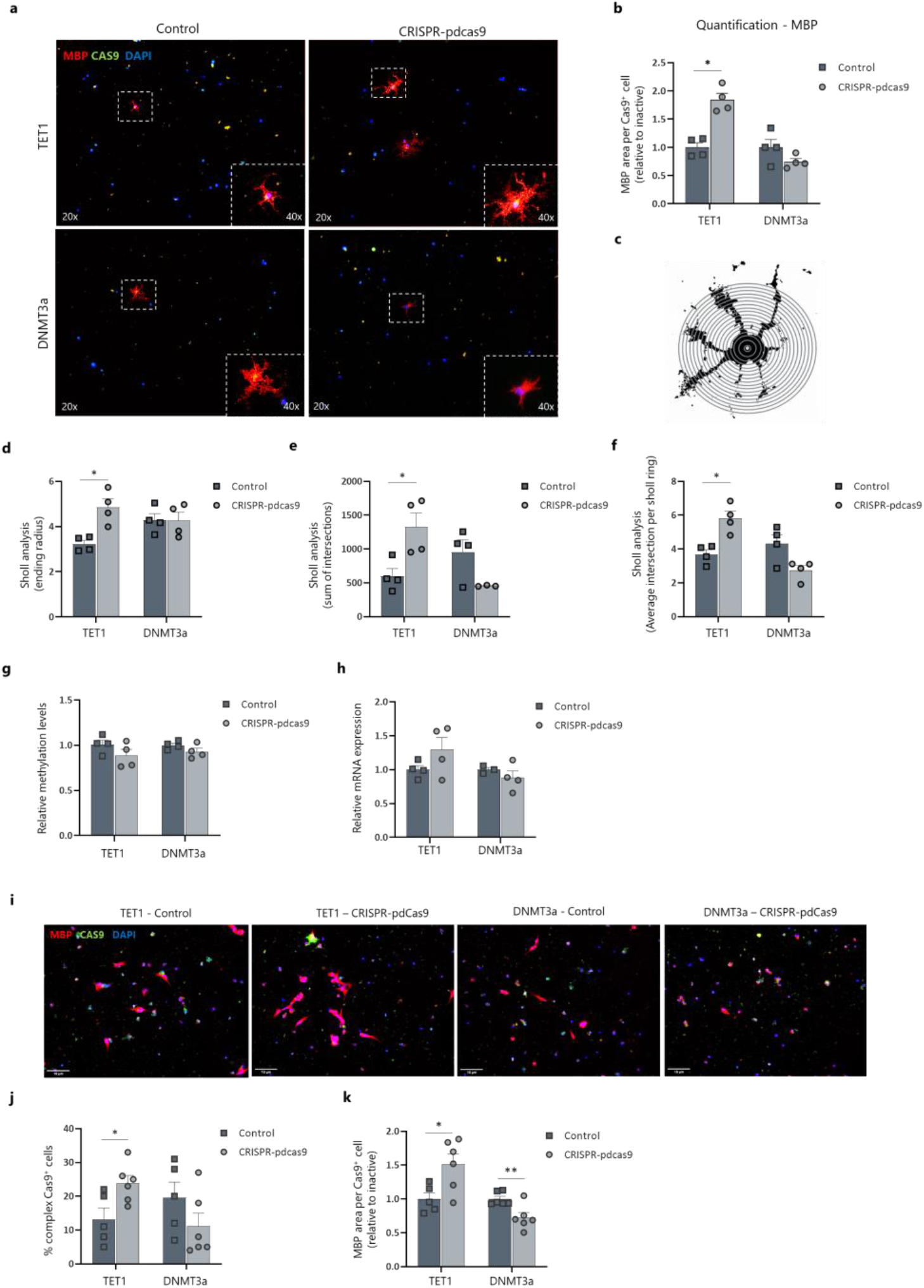
Epigenetic editing of the *MBP* promotor region in human iPSC-derived oligodendrocytes and a human oligodendroglioma (HOG) cell line influences the differentiation capacity. Human iPSC-derived oligodendrocytes and human oligodendroglioma cells were transfected with either a CRISPR-pdCas9-DNMT3a or CRISPR-pdCas9-TET1 vector to methylate or demethylate the promotor region of the *MBP* gene. Inactive constructs harbouring a catalytical inactive DNMT3a or TET1 were used as control. **a** Representative images of transfected human iPSC-derived oligodendrocytes. **b** Quantification (MBP fluorescence area) of transfected human iPSC-derived oligodendrocytes show an effect on MBP protein expression after epigenetic editing (n=4, Wilcoxon test). **c-f** Representation and quantification of the Sholl analysis of transfected iPSC-derived oligodendrocytes (n=4, Wilcoxon test). **g** Methylation analysis of the *MBP* promotor region after epigenetic editing of the *MBP* promotor (n=4). **h** Gene expression analysis showed a tendency towards an altered expression profile of *MBP* after targeted (de)methylation. Data are corrected for the most stable housekeeping genes (RPL13a and TBP), n=4. **i-k** Representative images and quantification (complexity and MBP fluorescence area) of transfected human oligodendroglioma cells also show an impact of epigenetic editing on cellular behaviour (n=6, unpaired t-test). Data are represented as mean ±SEM, *p<0.05, **p<0.01.

## Discussion

In the current study, we investigated the transcriptomic and epigenomic profile of chronically demyelinated lesions and the surrounding NAWM from nine donors, with the final goal of understanding the molecular mechanisms underlying the hampered differentiation capacity of OPCs within the MS lesion microenvironment. We found 641 genes to be differentially expressed between lesions and NAWM. Subsequent methylation analysis on this gene set revealed a total of 8,336 CpGs located on 512 different genes displaying differential methylation between lesions and NAWM. Gene ontology analysis revealed enriched clusters of genes related to glial cell development and myelination. We then further explored *MBP*, the gene with the highest number of DMPs within the promotor region among these clusters. This gene displayed decreased expression as well as hypermethylation in lesions. Cell-specific validation of *MBP* methylation in lesion-derived OPCs revealed a similar hypermethylated profile compared to NAWM-derived OPCs. Finally, we functionally validated the influence of *MBP* methylation on oligodendrocyte differentiation by means of epigenetic editing.

The involvement of DNA methylation in oligodendrocyte differentiation has been investigated previously by us and other researchers (6, 13, 14, 32). Using rodent-derived OPCs or mouse models for MS, it has been shown that the presence of DNA methylation enzymes, such as DNMT1 and DNMT3a, is crucial for oligodendrocyte differentiation during development and remyelination (14, 15). Furthermore, we have recently established that the myelin regulatory pathway, *Id2* and *Id4* in particular, is under epigenetic control during physiological OPC differentiation (13). Yet, the direct impact of DNA methylation in relation to remyelination failure in MS remained to be investigated. One of the first studies to use an epigenome-wide approach investigated methylomic alterations within NAWM brain samples of MS patients and compared them to matched non-neurological white matter control samples (16). Pathology-free MS samples show differentially methylated regions within genes related to oligodendrocyte development and survival. In line with this notion, recent studies by Kular *et al*. showed specific DNA methylation profiles of neuronal and glial cells isolated from the NAWM of post-mortem MS brains (17, 18). As for lesions, one study has investigated the difference in the methylation patterns between demyelinated and intact hippocampi of progressive MS patients using the Illumina Methylation 450K array. Chomyk et al. elegantly highlighted several DMPs related to neuronal survival and memory function yet did not reveal any methylation changes related to oligodendrocyte biology (33). Altogether, it is evident that the DNA methylation is affected in MS post-mortem brain tissue, but how this relates to the block on OPC differentiation in chronically demyelinated lesions has remained unclear up to now.

In the present study, we aimed to investigate the methylomic signature of chronically demyelinated MS lesions in order to understand the direct contribution of DNA methylation to the hampered differentiation state of OPCs within these lesions. One of the main strengths of this study is the unique within-comparison between lesions and their surrounding NAWM isolated from each patient. This setup increased our statistical power and allowed us to investigate DNA methylation changes specifically related to the lesion microenvironment, where OPC differentiation is hampered. Furthermore, we examined both transcription and DNA methylation in these samples, allowing us to directly correlate our transcriptional data to the methylation profile of these genes.

Our GO analysis based on genes that displayed both differential expression and methylation, as well as a significant correlation between these two features, revealed two main clusters, i.e., ‘cytoskeleton organisation’ and ‘glial cell development and myelination’. Genes within these clusters ranged from important myelin genes (*MBP, MAG*) and genes that regulate myelin formation (*CNTN2, LPAR1*) or OPC differentiation (*PARD3, BCAS1*), to genes important for lipid metabolism (*UGT8, ABCA2*) (34–39). Intriguingly, we found *MBP* to contain both the highest number of DMPs overall and the highest number of DMPs located within the promotor region and on CpG islands or shores. Moreover, all the probes within the promotor region of *MBP* were consistently hypermethylated in lesions compared to the surrounding NAWM. One could advocate that the *MBP* gene would be an obvious suspect to be altered within a demyelinated lesion. Interestingly, *MBP* has also been shown to be hypermethylated in NAWM samples of MS patients compared to non-neurological controls (16). These findings suggest a possible step-wise methylation change in the *MBP* gene, already initiated in NAWM regions and becoming more pronounced in the actual lesion site, where myelin damage has already occurred. Moreover, *MBP* has also been shown to be differentially methylated in other neurodegenerative diseases with white matter pathology, such as Alzheimer’s disease (AD). A recent meta-analysis, which combined data from six independent brain AD methylation studies (n=1,453 individuals), investigated the methylation profile of 485,000 CpG sites, of which one of the differentially methylated CpG sites in the prefrontal cortex reaching genome-wide significance was located within the *MBP* gene (40). Altogether, this emphasises the importance of DNA methylation in the regulation of *MBP* expression and its susceptibility to changes during disease.

The observed hypermethylated profiles of *MBP* within NAWM and MS lesions represent interesting independent observations, yet they could potentially be explained by differences in cellular composition between patients and controls and between lesions and NAWM, respectively (16). We however hypothesized that the epigenetic block on *MBP* was present in OPCs within lesions, thereby inhibiting their differentiation into myelin-forming oligodendrocytes. As such, we laser-captured OPCs both from lesions as NAWM samples and performed bisulfite pyrosequencing of the *MBP* promotor region. Altogether, these results indicate that the *MBP* promotor becomes hypermethylated in OPCs located within the lesion microenvironment, possibly preventing them from differentiating into mature oligodendrocytes. Indeed, *in vitro* experiments using OPCs isolated from *shiverer* mutant mice, showed that the lack of *MBP* does not impact their proliferative capacity, but prevents their differentiation process towards mature oligodendrocytes (41). While the absence of other myelin proteins, such as PLP and CNP, does not affect CNS myelination, the lack of functional *MBP* results in severe myelination deficits (42, 43). Next to its crucial role in myelin compaction, MBP also facilitates glial cytoskeleton assembly during oligodendrocyte differentiation. OPC undergo major morphological changes during their differentiation into oligodendrocytes, which requires microfilament and microtubule network remodelling (44). MBP has been show to polymerise and bundle actin filaments and microtubules, cross-link them to each other, and facilitate their binding to the lipid membrane (45–48). In line, oligodendrocyte cultures derived from *shiverer* mice fail to form proper myelin sheaths due to abnormally assembled microtubule and actin-based structures, thereby emphasising the essential role of MBP during oligodendrocyte differentiation (44, 49).

Our observations regarding *MBP* methylation in MS lesions and lesion-derived OPCs are novel, yet remain correlational. As we have suggested previously, it is important to investigate the potential cause-and-effect relationship between epigenetic signatures and functional read-outs, such as oligodendrocyte differentiation (31). Over the past years, epigenetic editing, using CRISPR-dCas9 engineered systems, has proven to be a powerful tool to provide evidence of functional consequences of epigenetic changes at specific loci (50). In the present study, we made use of both a CRISPR-dCas9-TET1 and a CRISPR-dCas9-DNMT3a vector to target and demethylate or methylate, respectively, the *MBP* promotor region with the final aim of assessing the influence on oligodendrocyte differentiation capacity (51, 52). We used two cell culture models, i.e. human iPSC-derived oligodendrocytes and HOG cells, which we transfected with the epigenetic editing vector to target the *MBP* gene, and assessed the effects on MBP protein expression and cellular morphology. As a control for transfection and steric hindrance, we transfected cells with a catalytic inactive version of the vectors that are unable to (de)methylate. Interestingly, we observed significant functional effects after targeted demethylation of the *MBP* gene, resulting in higher MBP expression and a more differentiated cellular morphology, both in iPSC-derived oligodendrocytes and in the HOG cell line. Targeted methylation showed less pronounced effects, yet did reveal a consistent trend towards reduced MBP protein expression and lower cellular complexity. A possible explanation for this could be that the baseline default methylation status of both cell culture types already levelled around 80%, leaving little room for effects of additional (hyper)methylation by the CRISPR-dCas9-DNMT3a vector. A limitation of the CRISPR-dCas9-based epigenetic editing tool, is the possible off-target effects due to undesired binding to other genomic regions. The endonuclease inactive CRISPR-dCas9-based epigenetic editing tool acts mainly as a cargo protein to guide the effector domains to the desired region, and is therefore more tolerant to mismatches between the sgRNA and the DNA loci (53). As we cannot completely rule out possible off-target events, we designed our sgRNA in such way that the predicted off-target sites were located in non-coding genomic regions.

Collectively, our data demonstrate strong differences in DNA methylation between chronically demyelinated MS lesions and the NAWM, which furthermore correlate with the expression profile of the corresponding DEGs. We identified an epigenetic block on *MBP* within OPCs located in the lesions and showed that this could have a major impact on the differentiation capacity of these cells. Notably, more than 8,000 CpG sites displayed differential methylation within MS lesions, with numerous of them potentially impacting upon cellular behaviour within the lesion site. It is therefore important to further characterise MS-associated epigenetic signatures, preferably in a cell type-specific manner, in order to fully understand the contribution of DNA methylation to remyelination failure in progressive MS stages. Which specific molecules and factors within the microenvironment of demyelinated lesions drive the observed epigenetic changes remains to be elucidated. Our study represents a starting point for important research regarding DNA methylation signatures in chronic MS lesions with the final aim to discover new targets to restore the remyelination capacity during progressive MS.

