## Supplementary information for "From methylation to myelination: epigenomic and transcriptomic profiling of chronic inactive demyelinated multiple sclerosis lesions"

**Supplementary Table 1: Guide RNA targeting MBP**

| | Forward oligo (5’-3’) | Reverse oligo (5’-3’) | Off-target score |
|---|---|---|---|
| *MBP* sgRNA | ACTGACTCCAAGCGCACAG | CTGTGCGCTTGGAGTCAGTC | 63 |

**Supplementary Table 2: Pyrosequencing primer list**

| Target gene | Forward primer (5’-3’) | Reverse primer (5’-3’) | Sequencing primer (5’-3’) | Genomic coordinates | Number of CpGs covered | Reference genome |
|---|---|---|---|---|---|---|
| *MBP* | GTTTGGTAGGATGTTTATTTAGTTGA | TCTATAACCCCATCACATCCAAACTCTC | GGATGTTTATTTAGTTGATTTAGG | chr18: 77016996-77017182 | 4 | GRCh37 (hg19) |

**Supplementary Table 3: qPCR primer list**

| | Forward oligo (5’-3’) | Reverse oligo (5’-3’) |
|---|---|---|
| *MBP* human | AAGACAGGCCCTCTGAGTCC | GGAGGGTCTCTTCTGTGACG |
| *TBP* human | TATAATCCCAAGCGGTTTGC | GCTGGAAAACCCAACTTCTG |
| *RPL13a* human | AAGTTGAAGTACCTGGCTTTCC | GCCGTCAAACACCTTGAGAC |

**Supplementary Table 4: Antibody list**

| Antigen | Company (reference number) | Dilution |
|---|---|---|
| MBP (iPSC-oligo’s) | Merck (MAB386) | 1:500 |
| MBP (HOG cells) | MilliporeSigma (AB980) | 1:500 |
| CAS9 | Merck (MAC133) | 1:1000 |
| Goat anti-mouse IgG | Life Technologies (AB_2534069) | 1:600 |
| Goat anti-rat IgG | Life Technologies (A21042) | 1:600 |
| Goat anti-rabbit IgG | Life Technologies (A27039) | 1:600 |

**Supplementary Table 5: Predicted off-target sites of the sgRNA**

| Sequence | PAM | Gene | Locus |
|---|---|---|---|
| GACTGACTCCAAGCGCACAG | CGG | MBP | chr18:-77017196 |
| TTCTGACTCCAAGCACACAG | AGG | / | chr20:-62149089 |
| GAAGGACGCCAAGCACACAG | TAG | / | chr19:+10269517 |
| GACT-ACCCCAAGCACACAG | AAG | / | chr5:+153455673 |
| TATTGACCCCAAGCACACAG | TAG | / | chr17:-39911657 |
| GGCTGAATCCAAGCACACAA | GAG | / | chr8:-40385002 |
| GACAGAGCTCCAAGCGCACAA | AAG | / | chr10:-84054755 |
| GGCTGAGGCCAAACGCACAG | CAG | / | chr8:+142798613 |
| GACTAACGCCAAGC-CACAG | AGG | / | chr10:-117736779 |
| GACT-ACTCCAACCACACAG | GAG | / | chr11:-130576698 |
| GACTTA-TCCAAGCGAACAG | TGG | / | chr19:+33712560 |
| AACTCACACCAACCGCACAG | AAG | / | chr8:+13309745 |
| CCCTGACCCCAAGCCCACAG | TAG | / | chrX:-2692829 |
